# Differential Effects of Meditation States and Traits on the Neural Mechanisms of Pain Processing

**DOI:** 10.1101/2025.05.20.655049

**Authors:** Vasil Kolev, Peter Malinowski, Antonino Raffone, Valentina Nicolardi, Luca Simione, Salvatore M. Aglioti, Juliana Yordanova

## Abstract

**Objectives:** The main objective of the present study was to explore the effects of different types of meditation on the neurophysiologic mechanisms of pain processing.

**Methods:** EEG responses to electric median nerve stimulation were recorded in short-term and long-term meditators (STM, LTM) during rest and three forms of meditation engaging attentional and affective regulation in different ways: focused attention meditation (FAM), open monitoring meditation (OMM) and loving kindness meditation (LKM). EEG responses were analysed in the time- and time-frequency domains to compute local components, and temporal and spatial synchronizations of multi-spectral pain-related oscillations (PROs) in order to characterize bottom-up processes, pro-active modulation of cortical excitability, cognitive/affective appraisal, and the connectivity of performance monitoring (fronto-medial) and attentional (fronto-parietal) networks during pain processing.

**Results:** STM manifested a significant decrease in the connectedness of the fronto-medial theta-alpha network and a significant reduction of the P3b during LKM. In contrast, changes in LTM were observed during FAM and OMM. They were characterized by pre-stimulus alpha increase at somatosensory areas, and modulations of fronto-medial and fronto-parietal theta-alpha synchronizations.

**Conclusions:** Different meditation states do not influence bottom-up sensory pain processing. However, they significantly alter cognitive/affective pain mechanisms in state- and trait-dependent ways. In novice meditators, a positive emotional disposition during meditation can suppress the distribution and cognitive/affective appraisal of nociceptive signals. In expert meditators, effects of meditation states on pain processing are critically guided by advanced control of internal attention leading to fine-tuned involvement and functional segregation of cognitive control and attention networks.

## 1. Introduction

Meditation states and traits have been consistently associated with a decrease in perceived subjective pain. These positive effects are suggested to result mainly from altered processing of emotional and attention-related components of pain (Bakhshani et al., 2016; Brefczynski-Lewis et al., 2007; Grant & Rainville, 2009; Grossman et al., 2007; Kasai et al., 2017; Morone et al., 2008; Nicolardi et al., 2022; Nielsen & Kaszniak, 2006; Schmidt et al., 2011; Su et al., 2016) though the precise neural mechanisms remain debatable. One neurophysiological model posits that major effects of meditation on pain rely on ***reactive cognitive control***, whereby noxious stimuli are sensed as really painful but the unpleasantness/affect (i.e., the emotional/cognitive appraisal) is reduced or does not occur (Gard et al., 2012; Grant et al., 2011; Jinich-Diamant et al., 2020; Perlman et al., 2010; Zeidan & Vago, 2016). Thus, the sensory experience of pain is separated from the subsequent negative evaluation (Zorn et al., 2020, 2021). An alternative model posits that ***pro-active cognitive mechanisms*** are crucial for pain processing in meditation (Zeidan et al., 2011, 2015). According to this model, effortful cognitive processes mediated by executive attention downregulate pain signals in the thalamus and transform ascending nociceptive information from painful to innocuous (sensory) thus avoiding the need for further extensive cognitive/affective reappraisal (Zeidan et al., 2011; Jinich-Diamant et al., 2020).

However, a great variety of meditation practices exist (Chiesa & Malinowski, 2011; Davidson & Dahl, 2017; Lutz et al., 2015). As outlined below, a neurocognitive classification of meditation practices from different traditions has identified three major meditation types with specific neurocognitive characteristics (Cahn & Polich, 2006; Dahl et al., 2015; Lutz et al., 2008; Raffone et al., 2014, 2019; Vago & Silbersweig, 2012). As representing specific brain states, they may modulate pain sensation and related neural processes in distinct ways (Wiech, 2016).

*Focused Attention Meditation* (FAM) entails voluntary focusing attention on a chosen object in a sustained fashion. The attentional focus in FAM can be on physical objects in the external world, bodily sensations, such as the breath, but also on mental objects, including thoughts, emotions or imagined visual forms. Focusing and sustaining attention on a single object requires maintaining object representation, controlling the focus of attention, detecting distraction, disengaging attention from the source of distraction, and (re)directing and engaging attention to the intended object (Lutz et al., 2008, 2015; Malinowski, 2013). Hence, specific neural systems associated with selective attention, sustained attention and conflict monitoring are involved in inducing and maintaining the state of FAM (Lutz et al., 2008; Hasenkamp et al., 2012; Yordanova et al., 2021). Upon pain stimulation in FAM, focusing attention on a single object different from pain is expected to re-allocate attention away from pain stimuli.

*Open Monitoring Meditation* (OMM) entails moment-to-moment monitoring of the contents of experience, mental processes or any object appearing in the present moment in a non-reactive way, by avoiding explicit attentional selection or focus on any specific object (Lutz et al., 2008). OMM requires awareness and meta-awareness of momentary sensations, thoughts and feelings, an ability closely linked to mindfulness (Isbel & Summers, 2017; Malinowski, 2013; Raffone et al., 2019). In contrast to FAM requiring the narrowest attention focus, OMM regulates attention with the widest possible aperture (Lutz et al., 2015; Dahl et al., 2015). Hence, the induction and maintenance of OMM relies on brain systems associated mainly with divided attention, monitoring, awareness, and attention/mind wandering regulation (Isbel & Summers, 2017; Malinowski, 2013; Raffone & Srinivasan, 2009). Upon pain stimulation in OMM, the imposed enhanced monitoring and awareness of all external and internal events would include pain, but limited emotional reactivity and cognitive evaluation of pain stimulus is expected.

*Loving Kindness Meditation* (LKM) is another major meditation style, which refers to the cultivation and maintenance of positive affective mental states associated with caring feelings, attitudes and intentions, acceptance, and motivation toward wellbeing of self and others, with equanimity (Dahl et al., 2015; Lutz et al., 2008). In contrast to FAM and OMM, LKM does not emphasize attention and meta-cognitive awareness although it also requires a certain degree of top-down regulation of mind wandering, attentional control and monitoring of meditation-relevant representations (Dahl et al., 2015; Raffone et al., 2014, 2019). Upon pain stimulation, LKM may primarily affect the cognitive/affective appraisal of pain stimuli.

Since these three forms of meditation may plausibly involve specific brain states by engaging attention and emotion systems in distinct ways (e.g., Lutz et al., 2008; Yordanova et al., 2020, 2021), one aim of the present study was to characterize the neurophysiologic pain mechanisms during FAM, OMM, and LKM. Another aim was to assess if meditation-specific pain processes might differ between practitioners with short or long meditation practice. For these aims, in the present study, electroencephalographic (EEG) responses to pain were recorded during rest and three meditation conditions – FAM, OMM and LKM – in two groups of meditators - novice meditators with little meditation experience, here termed short-term meditators (STM) and highly-experienced meditation experts, here termed long-term meditators (LTM). A set of neuroelectric parameters was selected for analysis to reflect bottom-up and top-down pro-active and reactive pain processes:

### (A) Bottom-up somatosensory processes

EEG pain research has demonstrated that evoked pain-related potentials (PRPs) and phase-locked oscillatory alpha-to-gamma responses to noxious stimuli at primary somatosensory (S1) and secondary somatosensory/insular (S2-IC) regions are predominantly associated with bottom-up processes (Babiloni et al., 2002; Hauck et al., 2015; Nickel et al., 2022; Strube et al., 2021; Tiemann et al., 2015). In the present study, local time domain PRP components and the temporal phase-synchronization of multi-spectral pain-related oscillations (PROs) were analysed to reflect the effects of specific meditation states on bottom-up pain processes.

### (B) Communication of pain information across primary, secondary and associative pain-processing regions

In contrast to bottom-up processes, top-down influences on pain processing have been mainly associated with spatial synchronization of alpha and gamma oscillatory networks after painful stimulation (Bott et al., 2023). Further, it has been shown that the spatial synchronization of S1 and S2-IC regions following a pain stimulus is sensitive to meditation expertise (Yordanova et al., 2025). To study if specific meditation states are associated with distinct communication patterns across key regions of pain processing (Peyron et al., 2000), the spatial synchronization of theta-to-gamma oscillatory networks between S1, S2-IC and fronto-medial (FM)/ACC regions was analysed.

### (C) Pro-active processes of pain expectancy

Pro-active top-down influences on somatosensory processing have been correlated with ongoing oscillatory activity from alpha and beta frequency ranges preceding event delivery (Babiloni et al., 2006; Pfurtscheller & Lopes da Silva, 1999; van Ede et al., 2010, 2011). Specifically, attention orientation to upcoming painful stimuli has been consistently verified by a pronounced alpha desynchronization at the contra-lateral S1 and S2 (Babiloni et al., 2003, 2004, 2006; Peng et al., 2015; Ploner et al., 2006a, 2006b, 2017). In the present study, to explore pro-active preparatory attention to pain stimulus in different meditation states, alpha and beta activities preceding pain stimulus delivery were analysed.

### (D) Reactive top-down processes of cognitive and emotional appraisal of pain information

Extensive research has provided consistent evidence for the association between the late parietal P3b component of event-related potentials and the amount of attention allocated to cognitive stimulus evaluation (Polich, 2007). P3b has been identified as a reliable index of involuntary attentional shifts and emotional processing of nociceptive events (Legrain et al., 2002, 2009, 2012). In the present study, the P3b component of time-domain PRPs was analysed to explore whether meditation might influence the reactive top-down processes of cognitive and emotional appraisal of pain information, and if such effects differ between FAM, OMM, and LKM.

### (E) Activation of cognitive control networks

Previously, significantly enhanced theta (4-8 Hz) oscillations have been observed at medial-frontal electrodes (centered on FCz) in different sensorimotor conditions in relation to a variety of executive and cognitive control functions - conflict processing, detection of errors, inhibition, performance monitoring, amount of cognitive control, and behavioural re-adjustment (Cavanagh & Frank, 2014; Cavanagh et al., 2009; Cohen & Donner, 2013; Cohen & van Gaal, 2013; Cohen, 2011, 2014; Fusco et al., 2018, 2022; Kolev et al., 2009, 2024; Nigbur et al., 2012; Yordanova et al., 2004, 2020, 2024). It has been posited that a theta network “hub” in the medial fronto-central cortex serves to coordinate behaviour in different contexts thus supporting various executive functions during cognitive control (Cohen, 2011, 2014; Duprez et al., 2020). Previous studies in no-pain resting states have revealed a substantial reorganization of the fronto-medial theta “hub” in experienced meditators as compared to novices (Jo et al., 2017; Yordanova et al., 2021). Thus, the connectedness of the fronto-medial area was studied here to explore if the maintenance of different meditative brain states might rely on specific activations of the medio-frontal cognitive network.

### (F) Activation of attentional networks

Fronto-parietal (FP) networks have been recognized as the neural substrate of focused attention and attention re-allocation (Corbetta & Shulman, 2011; Vossel et al, 2014). Specifically, the dorsal FP networks are involved in the control of spatial and feature-based attention and in stimulus-response mapping, while the right-lateralized ventral FP network is linked to reorienting of attention to unexpected but behaviourally relevant events (Chica et al., 2013; Corbetta & Shulman, 2002; Vossel et al., 2014). FP networks also sub-serve attention-based conscious perception (Chica et al., 2013; Dehaene & Changeux, 2011; Dehaene et al., 1998; Rees, 2013), cognitive control and working memory (Bressler & Menon, 2010; Egner et al., 2008; Menon, 2013; Rottschy et al., 2012, 2013). Major operating frequencies of FP networks have been established mainly for slow-frequency (theta) EEG ranges (Daitch et al., 2013; Sadaghiani & Kleinschmidt, 2016; Yordanova et al., 2017). Importantly, relevant activations of frontal and parietal regions have been observed during pain sensation (Peyron et al., 2000). However, the question of how FP networks contribute to pain processing during different meditation states has not been addressed systematically. In the present study, the FP synchronization of theta oscillations was computed to capture expected differences in the involvement of attentional networks during pain processing as a function of the specific meditation form.

A first hypothesis of the present study was that the neural mechanisms of pain processing would be specific for FAM, OMM, and LKM due to their different state-dependent and contextual top-down influences (Apkarian et al., 2005; Atlas, 2023; Kucyi & Davis, 2015; Peerdeman et al., 2016; Wiech, 2016). A second hypothesis was that the effects of specific meditation states on neural pain mechanisms would differ between STM and LTM groups. First, the two groups are expected to have dissimilar abilities in achieving and sustaining specific meditation states. Because long-term meditation practice is a dynamic process that is thought to entail a transition from initially effortful to effortless induction and maintenance of brain states, a considerable difference in the amount of the invested cognitive effort would accompany pain processing in the subjects from two groups while they sustain the specific states (Fell et al., 2010; Lutz et al., 2008). In addition, the pronounced neuroplastic modification of cognitive and pain-related networks in meditation experts (Fox et al., 2014; Guidotti et al., 2023; Lu et al., 2023; Tang et al., 2015; Yordanova et al., 2020, 2021) and the altered neural mechanisms of pain processing during resting state (Yordanova et al., 2025) strengthen the expectations for dissimilar neural reactions to pain in the two groups in specific meditation states.

## 2. Methods

The present study is based on a substantial data set used in previous reports (Nicolardi et al., 2022; Yordanova et al., 2025), but the results are new and have not been reported elsewhere.

### 2.1. Participants

The previously collected data from a total of 35 subjects were used, subdivided into a group of 19 long-term meditators (LTM; 3 females, mean age = 44.6 ± 10.9, mean number of years in monastery = 18 ± 12.7; mean lifetime duration of meditation practice 19358 hours (SE = 3164), range 900-50600 hours) and a group of 15 short-term meditators (STM; 6 females, mean age = 44.8 ± 8.2, with less than 250 hours of meditation experience). All participants were right-handed healthy volunteers, without a history of neurologic, psychiatric, chronic somatic, or other problems. The study had prior approval by the dedicated Research Ethics Committee at Sapienza University of Rome, Italy. All participants gave informed consent before participation, and the study was in accordance with the Declaration of Helsinki.

### 2.2. Tasks and Procedures

Neural responses to pain were elicited by applying painful electrical stimuli. The electrical stimuli were pulses generated by a monophasic constant current stimulator (STIM140, H.T.L. srl, Amaro, UD, Italy). Stimuli were delivered through two surface electrodes (diameter 6 mm, Ag-Cl, Electro-Cap International, Inc. Eaton, Ohio) placed 5 mm from each-other. The stimulation site was on the dorsal digital branch of the radial nerve, on the medial surface of the back of the left hand. The intensity range allowed by the instrument was between 2 and 50 mA.

Before experimental stimulation blocks, (1) the absolute pain threshold was determined, and (2) stimulus intensity calibration was performed. Using the ascending and descending method of limits (Säterö et al., 2000; Valentini et al., 2017), the absolute pain threshold of each participant was identified as the minimum intensity of a stimulus that was perceived as painful. During calibration, supra-threshold electric stimuli were delivered with a staircase procedure until the participant associated the same stimulus intensity with a moderate pain sensation in 50±10% of probes. Hence, individually adjusted stimuli with moderate intensity for each participant were used. During the experimental phase painful stimuli delivered as single events during conditions which the participants had to maintain in succession: a non-meditative resting state (REST), and meditation conditions (FAM, OMM, and LKM). The switch to each condition was signalled by means of an audio instruction. This sequence of meditative conditions was repeated 3 times, leading to a total of 12 blocks, with each block including 10 trials (a total of 30 trials for each condition).

The duration of each trial was approximately 10 s (9 to 13 s). The painful stimulus (electrical stimulus with 50 ms duration) was delivered randomly 4.5 to 8.5 s after beginning of the trial. One and a half second after stimulus delivery, participants were asked to rate (scale 1-100) three dimensions of subjective experience related to the nociceptive stimulation: pain, aversion, and identification (Nicolardi et al., 2022).

### 2.3. EEG recording and pre-processing

EEG recording and processing is presented in detail in Yordanova et al. (2025). Here, major analytic steps are summarized. EEG was recorded by a mobile wireless system (Cognionics; https://www.cognionics.net/mobile-128) using an electrode cap with 64 active Ag/AgCl electrodes located in accordance with the extended international 10/10 system and referenced to linked mastoids. Electrode impedances were kept below 10 kOhm and EEG signals were collected at a sampling rate of 500 Hz (resampled off-line to 250 Hz for data analysis).

EEG analysis was performed with Brain Vision Analyzer ver. 2.3.0 (Brain Products GmbH, Gilching, Germany). EEG was visually inspected for gross ocular and other artefacts at 64 channels. Contaminated trials were discarded along with EEG records exceeding ±100 µV. Bad channels were interpolated using topographic interpolation (Perrin et al., 1989). Slight horizontal and vertical eye movements preserved in the accepted trials were corrected by means of independent component analysis (ICA, Makeig et al., 1997). After artefact rejection, the mean number of artefact-free trials used for analysis was 27 for each condition (SD = 2.3, range 24-29).

To achieve a reference-free evaluation and control for volume conduction current source density (CSD) transform of the signals was performed (see, e.g., Nunez et al., 1997; Perrin et al., 1989 for details). All EEG epochs were analysed after a CSD transform of the signals.

### 2.4. Analysis of pain-related EEG responses

Pain EEG responses were analysed in the time- and time-frequency domains.

*Time-domain analysis* was performed by averaging 1.5 s long artefact-free pain-related EEG epochs, including 0.5 s before and 1 s after electric stimulus.

*Time-frequency (TF) analysis* of pain-related potentials was performed by means of a continuous wavelet transform (CWT) with Morlet wavelets as basis functions (Mallat, 1999).

Two types of epochs were analysed in the TF domain: (1) For slow-frequency PROs from delta, theta and alpha frequency bands, EEG epochs were 1.5 s long, including 0.5 s before and 1 s after electric stimulus, in the frequency range 0.5-25 Hz with a central frequency at 0.6 Hz intervals; (2) For fast-frequency PROs from beta and gamma frequency bands, EEG epochs were 0.85 s long, with 0.25 s before and 0.6 s after stimulus, in the frequency range 15-50 Hz with a central frequency at 1.75 Hz intervals.

### 2.5. Pain-related oscillations (PROs)

#### 2.5.1. Total power of PROs

Single trials were first transformed to the TF domain and then averaged. For each trial, the time-varying power for each frequency scale (total power, TOTP) was calculated by squaring the absolute value of the convolution of the signal with the complex wavelet. In the present study, TOTP was used for analysis of alpha and beta pre-stimulus activity. For statistical evaluation, TOTP was log10-transformed.

#### 2.5.2. Temporal synchronization of PROs: Phase-locking factor

The temporal phase synchronization across trials was analysed by means of the phase-locking factor (PLF, e.g., Lachaux et al., 1999; Tallon-Baudry et al., 1997; Yordanova et al., 2025). The values of PLF yield a number between 0 and 1 determining the degree of between-trial phase-locking, where 1 indicates perfect phase alignment across trials and values close to 0 reflect the highest phase variability. PLFs were computed for different TF scales at each time-point, each electrode, condition, and subject.

#### 2.5.3. Spatial synchronization of PROs: Phase-locking value

The phase-locking value (PLV) is measured between electrode channels and reflects the extent to which oscillation phase angle differences between electrodes are consistent over trials at each time/frequency point. PLV varies between one (constant phase difference) and zero (random phase difference). After excluding the edge electrodes prone to signal distortion and reducing the number of electrodes to 35, PLV was computed for each pair of electrodes, resulting in a total of 630 pairs for each subject. PLVs were computed for different TF scales at each time-point, each trial, condition and subject.

To identify regions with maximal connectivity during PRPs, the mean of all pairs connected with each single electrode was computed for each electrode. Following this procedure, a quantifier ‘regional PLV’ (R-PLV) was established. For R-PLV computation the selected 35 electrodes were used (F5, F3, Fz, F4, F6, FC5, FC3, FCz, FC4, FC6, C5, C3, Cz, C4, C6, CP5, CP3, CPz, CP4, CP6, P7, P5, P3, Pz, P4, P6, P8, PO7, PO3, POz, PO4, PO8, O1, Oz, and O2). R-PLV was computed for different TF scales at each time-point, each trial, electrode, condition and subject.

### 2.6. Measurements and Parameters

#### 2.6.1. Pain-related potentials (PRPs)

N1 PRP component was identified as the maximal negative peak within 130-250 ms after stimulus onset; P2 PRP component was identified as the maximal positive peak within 180-300 ms after stimulus onset; P3b was identified as the maximal positive peak with centro-parietal/parietal distribution within 300-500 ms after stimulus (Polich, 2007). Peak amplitude and latency of PRP components were measured against a baseline of 300 ms before stimulus.

#### 2.6.2. Pain-related oscillations (PROs)

As argued in Yordanova et al. (2025), PROs from 5 relevant frequency ranges were analysed: delta with central frequency *f_0_* = 3.12 Hz (1.5 – 4.5 Hz), theta-alpha with *f_0_* = 7.5 Hz (4.7 – 10.7 Hz), beta with *f_0_* = 18.14 Hz (15.8 – 22.9 Hz), gamma-1 with *f_0_* = 32.1 Hz (30.8 – 35.9 Hz), and gamma-2 with *f_0_* = 43.5 Hz (38.8 – 42.9 Hz). PLF and R-PLV were measured as the maximal value within defined epochs after stimulus for each frequency band, subject, and electrode as follows: delta (within 40-500 ms), theta-alpha (within 20-250 ms), beta (within 10-150 ms), gamma-1 (within 10-150 ms), gamma-2 (10-100 ms). The measures were baseline corrected by subtracting the mean value of a baseline of -300/-50 ms for slow and -200/-50 ms for fast frequency TF components.

#### 2.6.3. Pre-stimulus activity

Pre-stimulus activity was measured by computing the mean value of TOTP for alpha (*f_0_* = 7.5 and 10.1 Hz) within -450/-50 ms, and beta (-250/-50 ms) frequency bands. These were chosen for analysis of pre-stimulus activity as relevant for correlates of pro-active attention modulation (Peng et al., 2015; van Ede et al., 2010).

#### 2.6.4. Regions of interest

TF analyses were performed for sets of electrodes covering three regions of interest (ROIs): (1) contralateral primary somatosensory cortex (S1), (2) contralateral secondary somatosensory cortex including the insular cortex (S2-IC), and (3) frontal medial cortex including the anterior cingulate cortex (ACC) and supplementary motor areas (FM). Following the correspondence of electrode positions of the 10-10 system to these areas (Koessler et al., 2008; Scrivener & Reader, 2022), S1 included Cz, C4, CPz, CP4, S2-IC included C6, CP6, and P6, and FM included Fz and FCz electrodes (Babiloni et al., 2002; Bott et al., 2023; Nickel et al., 2022).

Figure 1 and Table 1 summarize the parameters analysed in the present study with respect to the objectives:

(A) To analyse bottom-up somatosensory processes: (1) time-domain PRP components N1 and P2 at S1 and S2-IC; (2) PLF of theta-alpha, beta, gamma-1 and gamma-2 PROs at S1, S2-IC, and FM – Fig. 1A.
(B) To analyse the communication of pain information across relevant cortical regions: PLV of theta-alpha, beta, gamma-1 and gamma-2 PROs of selected pairs of electrodes included in S1, S2-IC, and FM – Fig. 1B.
(C) To analyse pro-active processes of attention direction to or away from pain stimulus: pre-stimulus alpha and beta TOTP at S1, S2-IC, and FM – Fig. 1C.
(D) To analyse reactive top-down processes of cognitive/emotional appraisal of pain information: time-domain P3b component at centro-parietal electrodes CP3, CPz, and CP4 – Fig. 1D.
(E) To analyse the activation of the FM cognitive network: R-PLV of theta-alpha, beta, gamma-1 and gamma-2 oscillations at FM – Fig. 1E.
(F) To analyse the activation of attentional networks during pain processing: PLV of electrode pairs P5-F5, P3-F3, P4-F4, P6-F6. These pairs were selected to capture fronto-parietal synchronization of dorsal and ventral FP networks in the left and right hemisphere (Yordanova et al., 2017, 2021) – Fig. 1F.

**Figure 1.**
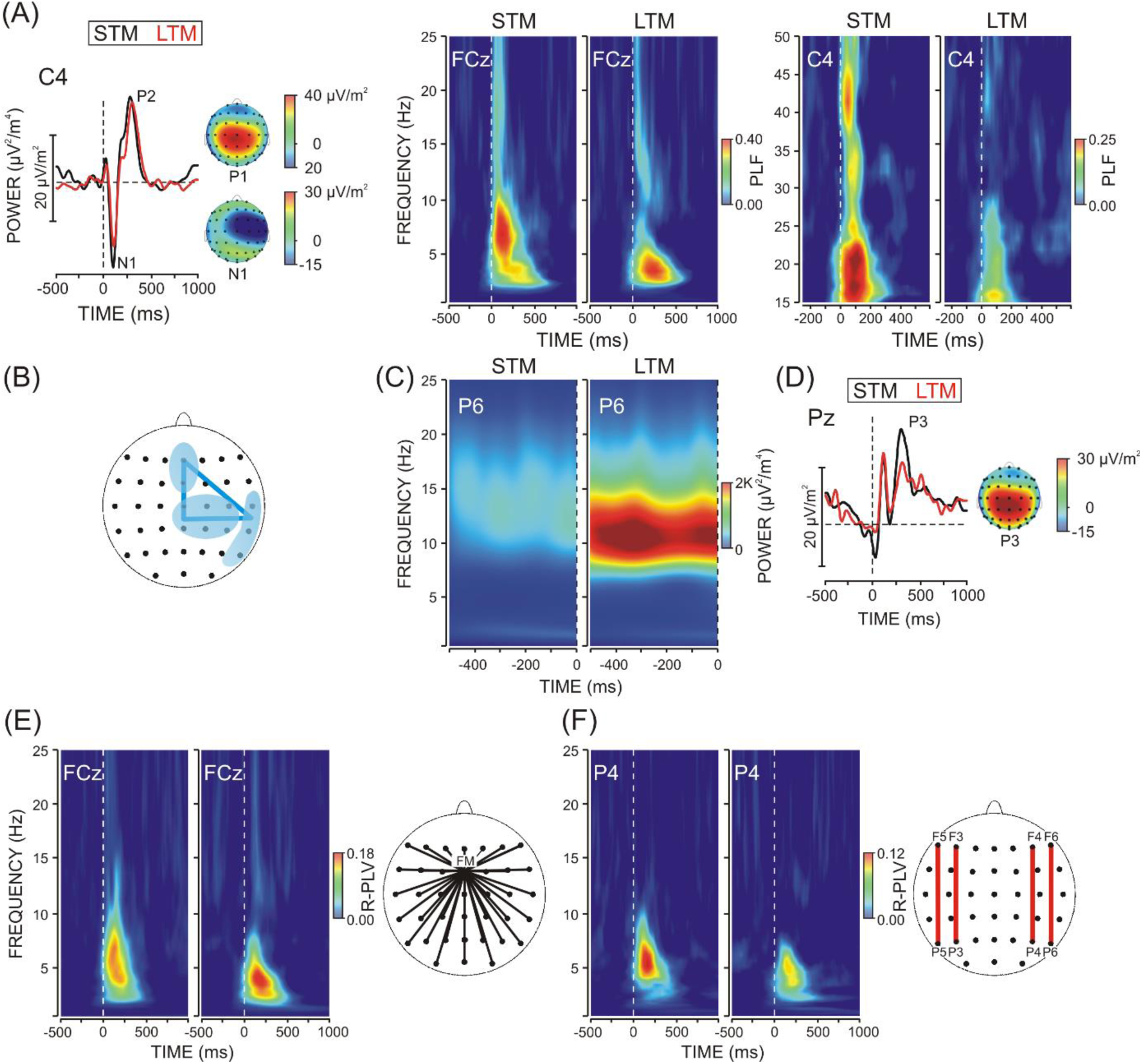
Neuroelectric parameters of pain processing in the two experimental groups – short-term meditators (STM) and long-term meditators (LTM). (A) Pain-related potentials (PRP) at S1 (C4 electrode) and topography maps of N1 and P2 PRP components (Left panel); Time-frequency plots of PLF for slow- and fast-frequency oscillations at relevant electrodes (FCz and C4) (Right Panel) during REST; (B) Schematic presentation of the connections across S1, S2-IC and FM regions analysed by means of inter-regional synchronization (PLV); (C) Pre-stimulus alpha activity (TOTP) during REST; (D) Waveform and topographic distribution of P3b PRP component of during REST; (E) Time-frequency plots of R-PLV during REST and a schematic presentation of the connectedness (R-PLV) of the fronto-medial region; (F) Time-frequency plots of PLV of fronto-parietal pairs during REST and a schematic presentation of 4 FP used for analysis. Stimulus occurrence at 0 ms.

**Table 1.**
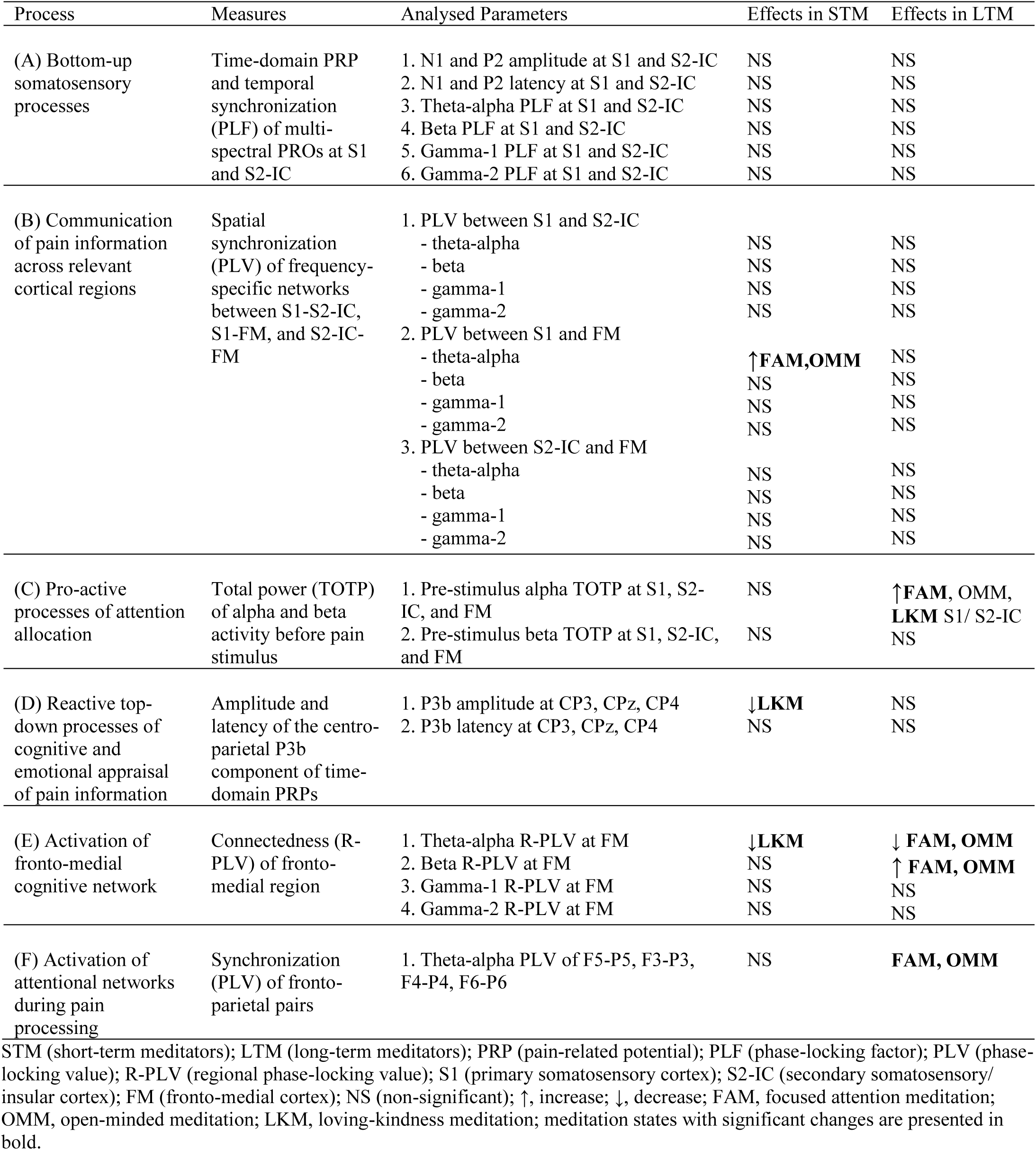
Summary of analysed neurophysiological processes, parameters, and effects.

### 2.7. Statistical analysis

In Yordanova et al. (2025), we demonstrated that PRPs and PRO parameters analysed here differ between STM and LTM already during REST, Therefore, the effects of meditation states were tested in each group separately. For each parameter, a repeated-measures ANOVA design was applied with a within-subjects variable Condition (REST vs. FAM vs. OMM vs. LKM). Planned contrasts were carried out to compare REST with each meditation state. Analyses were performed for each ROI (including electrodes relevant for S1, S2-IC, and FM) or selected PLV pairs, and each frequency-specific PLF/PLV/R-PLV/TOTP measure. For factors with more than two levels, the Greenhouse–Geisser correction was applied to the degrees of freedom (*df*). Original *df*, corrected *p*-values, and partial eta squared (ŋ^2^) are reported, along with mean group values ± standard error (SE) whenever relevant. To control for multiple testing effects, FDR correction was applied (Benjamini & Hochberg, 1995). For all analyses, only significant statistical outcomes are presented in detail in the results.

## 3. Results

### 3.1. Subjective pain indices

Figure 2 demonstrates that each of the parameters of subjective pain reflection (subjective pain intensity, aversion, and identification) decreased in LKM relative to REST in STM (Condition: subjective pain intensity, F(3/42) = 5.76, p = 0.013, ŋ^2^ = 0.307; aversion, F(3/42) = 10.45, p < 0.001, ŋ^2^ = 0.445; identification, F(3/42) = 8.02, p = 0.002, ŋ^2^ = 0.382; p < 0.05 for each contrast). The effects of meditation states on subjective pain measures were not significant in LTM (Condition; F(3/54) = 1.38, 3.07, p = 0.26, 0.08, ŋ^2^ = 0.112, 0.218) – Fig 2.

**Figure 2.**
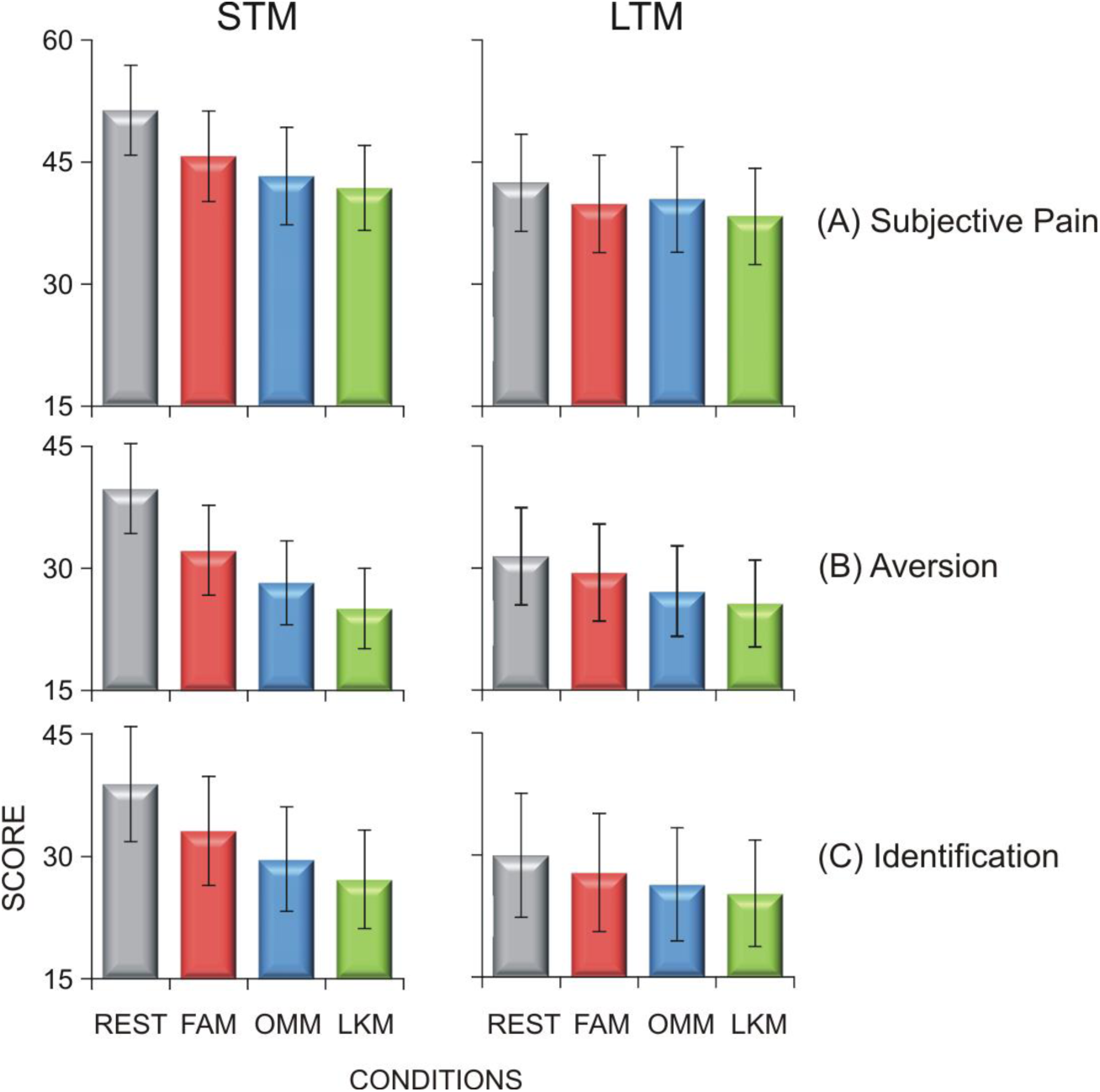
Subjective pain parameters: (A) subjective pain intensity, (B) aversion, and (C) identification in groups of short-term meditators (STM) and long-term meditators (LTM) in 4 conditions: REST, FAM, OMM, and LKM.

### 3.2. Neuroelectric parameters

#### 3.2.1. Bottom-up somatosensory processes

(Table 1, Fig. 3A). In none of the groups, was there any effect of meditation states on the parameters of local time-domain PRPs and PROs.

**Figure 3.**
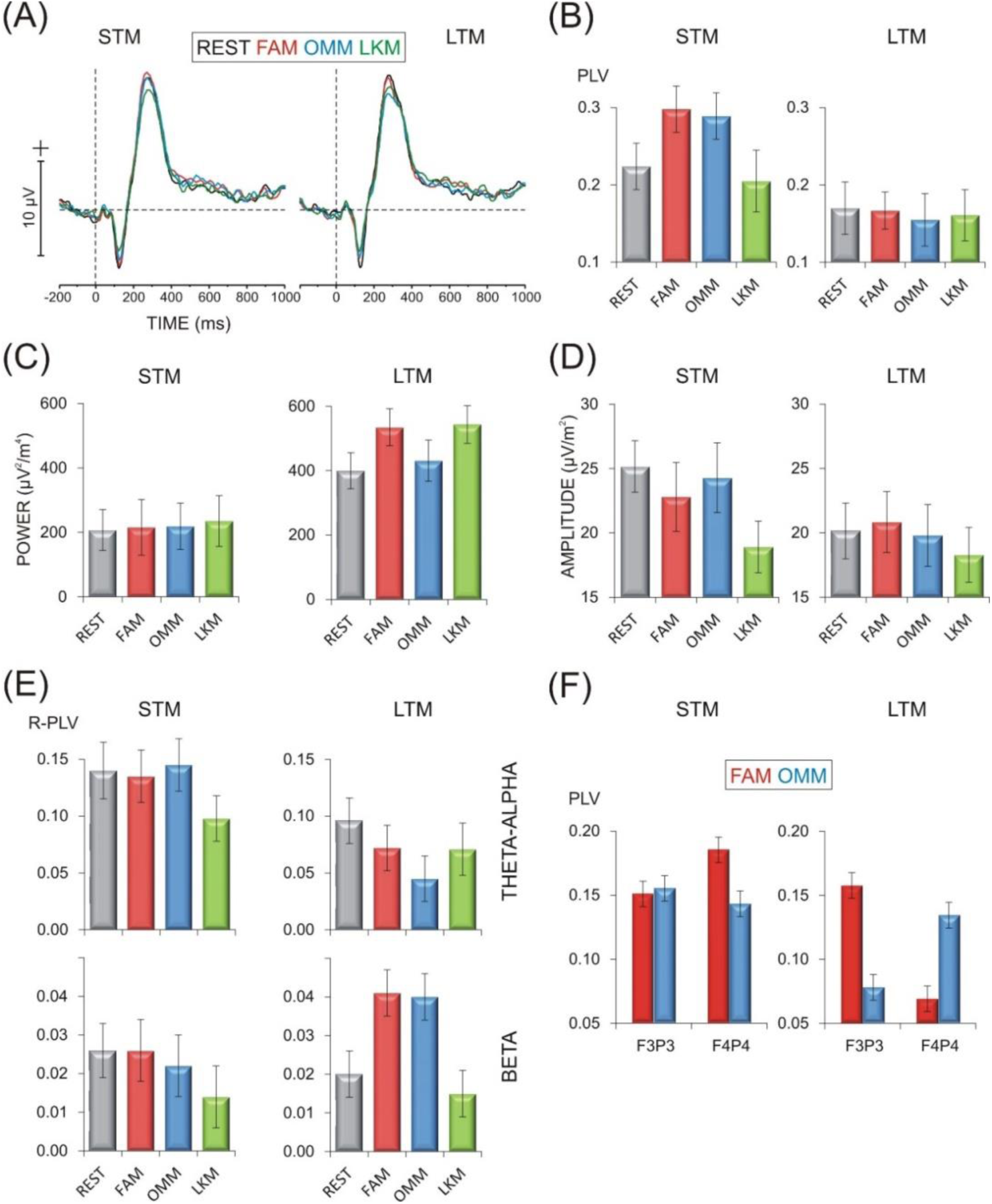
Group means ± SE of neuroelectric parameters of pain processing in 4 conditions (REST, FAM, OMM, and LKM) in two groups – short-term meditators (STM) and long-term meditators (LTM). (A) Grand average pain-related potentials (PRP) at S1 (C4 electrode); (B) Spatial synchronization (PLV) of theta-alpha networks between S1 and FM; (C) Pre-stimulus alpha activity (TOTP) at S1; (D) Amplitude of centro-parietal P3b component of PRPs; (E) Connectedness (R-PLV) of fronto-medial region supported by theta-alpha and beta oscillatory networks; (F) Synchronization (PLV) of fronto-parietal pairs F3-P3 and F4-P4 during FAM and OMM.

#### 3.2.2. Communication of pain information across cortical regions

(Table 1, Fig. 3B). No effects of meditation states were yielded for the synchronization between S1 and S2-IC for any of the frequency-specific oscillations in any of the two groups.

As demonstrated in Fig. 3B, in the group of STM, the synchronization between S1 and FM regions supported by theta-alpha networks significantly increased in FAM and OMM relative to REST (Condition, F(3/42) = 5.92, p = 0.01, ŋ^2^ = 0.297; REST vs. FAM, (F(1/14) = 15.02, p = 0.002, ŋ^2^ = 0.582; REST vs. OMM = 8.53, p = 0.01, ŋ^2^ = 0.389), whereas no difference in theta-alpha connections between S2-IC and FM was yielded. The synchronization between S1/S2-IC and FM supported by beta and gamma networks was not affected by meditation states in STM.

In the group of LTM, theta-alpha S1/S2-IC - FM connections were not modulated by meditation states (Fig. 3B), nor were there any reliable changes in the synchronization with FM of beta and gamma networks as a function of meditation states.

#### 3.2.3. Pro-active processes of attention direction

(Table 1, Fig. 3C). In the group of STM, pre-stimulus alpha activity did not vary with meditation states at any ROI: (Condition, F(3/54) < 0.681, p > 0.56, ŋ^2^ < 0.046).

In LTM, pre-stimulus alpha was significantly affected by meditation states at S1 (Condition, F(3/54) = 5.63, p = 0.006, ŋ^2^ = 0.238) and S2-IC (Condition, F(3/54) = 4.59, p = 0.03, ŋ^2^ = 0.203) by manifesting a significant increase in FAM (S1: F(1/18) = 6.34, p = 0.02, ŋ^2^ = 0.260; S2-IC: F(1/18) = 5.74, p = 0.03. ŋ^2^ = 0.242) and LKM (S1: F(1/18) = 15.46, p = 0.001. ŋ^2^ = 0.462; S2-IC: F(1/18) = 7.11, p = 0.01. ŋ^2^ = 0.283) as compared to REST. Meditation states in LTM were not associated with modulation of pre-stimulus alpha at the FM region. No effects of meditation states were detected for pre-stimulus beta activity at any ROI in any of the two groups (Condition, p > 0.3).

#### 3.2.4. Cognitive/emotional appraisal of pain information

(Table 1, Fig. 3D). Meditation states had an effect on P3b amplitude only in STM (Condition, F(3/42) = 5.53, p = 0.007, ŋ^2^ = 0.283) resulting from a significant P3b decrease in LKM as compared to all other states (F(1/14) = 4.83 – 10.86, p = 0.05-0.005, ŋ^2^ = 0.235 – 0.457). Interestingly, the maintenance of different meditation states in the group of LTM did not affect P3b amplitude (Condition, F(3/52) = 1.23, p = 0.3, ŋ^2^ = 0.064). No effects on P3b latency were observed.

#### 3.2.5. Fronto-medial networks

(Table 1, Fig. 3E). In STM, the FM theta-alpha synchronization was substantially reduced only during LKM as compared not only with REST but also with other meditation states, with no changes detected for FAM and OMM vs. REST (Condition, F(3/42) = 4.84, p = 0.009, ŋ^2^ = 0.257; LKM vs. REST, FAM, OMM, (F(1/14) = 5.44 – 21.44, p = 0.03 – 0.001, ŋ^2^ = 0.393 – 0.605).

In LTM, the effect of meditation states on FM theta-alpha synchronization also was significant (Condition, F(3/54) = 4.69, p = 0.01, ŋ^2^ = 0.207), but this was due to a significant reduction in all meditation states relative to REST, mostly expressed during OMM (REST vs. FAM, OMM, LKM, F(1/18) = 4.22, 22.56, 5.50, p = 0.05, 0.001, 0.03, ŋ^2^ = 0.152, 0.596, 0.234). Notably, opposite to theta-alpha, the FM beta synchronization was significantly enhanced during FAM and OMM in LTM (Condition, F(3/54) = 5.69, p = 0.008, ŋ^2^ = 0.307; REST vs. FAM, OMM, F(1/18) = 19.14, 20.05, p < 0.001, ŋ^2^ = 0.497, 0.502).

#### 3.2.6. Fronto-parietal networks

(Table 1, Fig. 3F). Four pairs of fronto-parietal (FP) electrodes were chosen to represent the dorsal and ventral attentional networks in the left and right hemisphere (F5-P5, F3-P3, F4-P4, and F6-P6). The FP synchronization was analysed in a Condition (4 levels) x Pairs (4 levels) repeated measures ANOVA design in each of the two groups.

In STM, meditation states were not associated with different synchronization of FP pairs (Condition, F(3/42) = 0.49, p = 0.7, ŋ^2^ = 0.037; Condition x Pair, F(9/126) = 1.33, p = 0.3, ŋ^2^ = 0.087).

However, in LTM, the synchronization of particular FP pairs was sensitive to specific meditation states (Condition x Pair, F(9/162) = 3.2, p = 0.05, ŋ^2^ = 0.187). Specifically, FAM and OMM produced lateralized modulations of theta-alpha FP synchronization. During FAM, an enhancement of F3-P3 synchronization was accompanied by a prominent decrease of F4-P4 synchronization (F(1/18) = 10.8, p = 0.004, ŋ^2^ = 0.375). Exactly the opposite modulations of F3-P3 and F4-P4 were observed during OMM, characterized by enhanced synchronization on the right and reduced synchronization on the left (F(1/18) = 5.54, p = 0.03, ŋ^2^ = 0.235). Of note, although non-significant, the opposite relations for the involvement of the right FP pair F4P4 in FAM and OMM was observed in STM.

## 4. Discussion

The present study aimed to characterize the effects of different meditation states on neurophysiologic mechanisms of pain processing. For that aim, three forms of meditation (FAM, OMM, and LKM) were explored, in which attentional and affective regulation were engaged in specific ways. Because extensive meditation practice cultivates advanced modes of awareness and attention/emotion regulation, two groups of meditation practitioners were studied, novices and experts. Several neurophysiologic indices of pain processes were analysed to characterize bottom-up and top-down mechanisms. It was expected that the states of meditation would affect neural mechanisms of pain processing in specific ways depending on (1) the form of meditation, and (b) meditation expertise.

### 4.1. Bottom-up mechanisms

One major result of the study is that the bottom-up mechanisms of pain processing do not differ significantly between meditation forms. We conclude this because the neural signatures that are sensitive to bottom-up processes (Bingel et al., 2004; Hauck et al., 2015; Nickel et al., 2022; Strube et al., 2021; Tiemann et al., 2015), namely the PRP components at contra-lateral S1 and S2-IC and the local temporal synchronization of multispectral pain-related oscillations, did not differ between meditation type or group.

However, it is important to note that our previous analyses during REST yielded differences between LTM and STM for the same parameters (Yordanova et al., 2025). These analyses revealed significantly lower phase-locked multispectral PROs at contra-lateral S1 and S2-IC regions in LTM than in STM, suggesting that a profound trait-related suppression of bottom-up pain mechanisms develops with meditation expertise. In this regard, if meditation is associated with a downregulation of ascending nociceptive signals in somatosensory areas (Brown & Jones, 2010; Jinich-Diamant et al., 2020, Zeidan & Vago, 2016; Zeidan et al., 2011), such sensory filtering may result primarily from neuroplastic network reorganization rather than from phasic brain states associated with specific meditation states.

### 4.2. Top-down mechanisms

In contrast to these bottom-up processes, the correlates of top-down mechanisms of pain were affected by different meditation states. Importantly, these differences also depended on meditation expertise, indicated by the differential effects of meditation states in novices and experts.

### 4.3. STM-specific effects

Specifically, STM manifested (1) enhanced theta-alpha synchronization of the S1-FM connections during FAM and OMM (Fig. 3B), (2) a significant decrease in the regional synchronization of the theta-alpha network centered at frontal midline regions during LKM (Fig. 3D), and (3) a significant reduction of P3b PRP component also during LKM (Fig. 3E; also see Table 1 for summaries). These results show that in STM, the most distinguished changes in pain processing are associated with LKM, i.e., the state imposing a shift toward positive emotional attitude, tone and reflection. Moreover, these mechanisms involved in STM during LKM appear to be specific as they were not sensitive to other meditation forms nor were they involved during LKM in experts. Likewise, the theta-alpha S1-FM connection that was only sensitive to attention-related meditation states (FAM and OMM) in STM, was not affected by any meditation form in experts.

### 4.4. LTM-specific effects

In contrast, in LTM, the most conspicuous state-related changes in top-down pain mechanisms emerged during FAM and OMM, i.e., meditation states that entail sustained attention regulation, in terms of a narrowed attentional focus for FAM and a maximally expanded attentional focus for OMM. Specifically, (1) pre-stimulus alpha increased at S1 and S2-IC, mostly in FAM (Fig. 3C), (2) the connectedness of the fronto-medial region supported by theta-alpha networks was suppressed in all meditation states, mostly in OMM (Fig. 3E top panels), (3) this suppressed theta-alpha fronto-medial synchronization was paralleled by increased beta synchronization of the fronto-medial region in FAM and OMM (Fig. 3E bottom panels), (4) there was a lateralized asymmetric synchronization of the fronto-parietal connections during FAM and OMM (Fig. 3F; also see Table 1 for a summary). These results reveal distinctive patterns in experienced meditators during FAM and OMM that did not occur in novices, pointing to highly specific effects of advanced attention/awareness control on neural pain processing.

We can now ask the question how these differential effects contribute to deepening our understanding of meditation-related pain processing in the brain:

### 4.5. Communication across regions of primary and higher-level pain processing

The synchronization between S1 and S2-IC can reflect a sequential (hierarchical) transmission of sensory pain information from S1 to S2-IC (Della Penna et al., 2004; Dockstader et al., 2010; Hadiwara et al., 2010) or a simultaneous co-activation of the two cortical regions (Garcia-Larrea et al., 2003). Although this communication may depend on focused attention (Dockstader et al., 2010; Hauck et al., 2007), no effects of meditation states were yielded here for any of the frequency-specific oscillations in either STM or LTM.

Only the synchronization between S1 and FM areas in the theta-alpha band in novices was altered during FAM and OMM. When interpreting these results we need to consider the fact that neural responses from the FM region capture activations of the anterior cingulate cortex (ACC), a key structure for integrative pain processing (rev. Peyron et al., 2000) that is involved in the affective and attentional components of pain sensation (Garcia-Larrea & Peyron, 2013; Garcia-Larrea et al., 2003; Peyron et al., 1999, 2000). A recent study in rats demonstrated that the sensory pain information encoded in the primary somatosensory cortex S1 is projected to a subset of neurons in the ACC, with increased ACC responses to noxious stimuli upon activation of the S1 axon terminals (Singh et al., 2020). In light of these findings, our results from STM suggest that the increased attention control (as in FAM and OMM) facilitates the transfer of noxious information from S1 to ACC. This increase, however, was not accompanied by higher subjective ratings of pain intensity or aversion (Fig. 2). Moreover, the projections from S1 to other pain regions (S2/insular) appear to be supported primarily by high-frequency (gamma) oscillations (e.g., Hagiwara et al., 2010), which were not selectively influenced by meditation states. Hence, we conclude that the enhanced theta-alpha S1-FM synchronization in STM may not merely index the transmission of specific noxious information from S1 to ACC. Instead, it may be a reflection of the involvement of ACC in domain-general integrative control systems operating in the theta frequency range (Cohen, 2011; Duprez et al., 2020). From this perspective, the results from STM imply that imposing a high load on attentional regulation in novices engages fronto-medial (ACC) control mechanisms whereby the communication with primary sensory regions is specifically emphasized. No such mechanisms were evident in LTM, suggesting that expert meditators do not strengthen connections between sensory and cognitive control regions upon intense attention regulation.

### 4.6. Pro-active processes of attention direction

Pre-stimulus alpha during meditation was only increased for LTM (Fig. 3C). This result extends our previous results from the same data set, which showed that by modulating ongoing alpha activity, expert meditators are able to pro-actively decrease the excitability of primary and secondary somatosensory cortices already during REST (Yordanova et al., 2025).

Although enhanced pre-stimulus alpha emerged in all meditation states, it was most pronounced in FAM and least pronounced in OMM. FAM involves a stable attentional focus to the relatively subtle sensation of breathing, which is likely to direct the attentional focus away from pain. Hence, we suggest that the additional amplification of pre-stimulus alpha at S1 and S2-IC during FAM corresponds to the re-direction of attention away from pain (Babiloni et al., 2006; Klimesch et al., 2007; Peng et al., 2015; Pfurtscheller & Lopes da Silva, 1999) that is central to FAM. In contrast, the wide attention aperture during OMM entails the detection and awareness of pain stimuli as inevitable ingredients of the entire information stream, which may have prevented further alpha amplification. These observations in LTM offer additional evidence regarding the ability of expert meditators to precisely tune the excitability of sensory regions in a pro-active manner through functionally distinctive influences from attentional systems on alpha oscillations (Hauck et al., 2007; Peng et al., 2015; Ploner et al., 2006a, 2006b). This may be critically associated with their capacity to sustain a stable and distinctive focus of internal attention (Lutz et al., 2008, 2015; Malinowski, 2013). Such a pro-active mechanism was not evident in novice meditators, potentially because they have not yet developed the same level of skill in sustaining specific modes of internal attention (Fell et al., 2010; Lutz et al., 2008). These results do not rule out that less experienced meditators are able sustain an attentional focus that is directed away from pain when attention is attracted to imperative or salient goals (Atlas, 2023; Legrain et al., 2005, 2011, 2013; Peerdeman et al., 2016). The important implication of the current results is that meditation experts, in contrast to novices, are able to deliberately inhibit/disinhibit sensory input at relevant cortical regions through alpha oscillations guided by advanced attention control.

### 4.7. Cognitive/emotional appraisal of pain information

The current result that P3b was reduced only in STM during LKM suggests a crucial role of affective states in novice meditators to restrain pain appraisal (Legrain et al., 2002, 2012). Hence, positive emotion states appear to diminish the cognitive/affective/motivational value of a pain stimulus leading to less re-allocated attention and less update in working memory of pain information (Polich, 2007). In support, this P3b decrease corresponded to lower subjective intensity, aversion and identification scores yielded only in novices during LKM (Fig. 2). In contrast, meditation forms did not influence substantially P3b in experienced meditators. This observation can be explained with generally reduced emotional involvement in LTM due to adopted equanimity attitude following long-term meditation training (Raffone & Srinivasan, 2009). Also, the present experimental design required subjective ratings after each pain stimulus, which might have preserved the cognitive evaluative component in LTM on the background of overall limited emotional processing (Singh et al., 2020).

### 4.8. Cognitive fronto-medial networks

The fronto-medial connectedness was analysed here to capture the activity of a critical cognitive hub including the ACC, which supports a variety of executive functions and integrates various information inputs to coordinate and synchronize behaviour (Cohen, 2011, 2014; Duprez et al., 2020). Although some sub-sections of the ACC are specifically responsible for integrating responses to pain (Wager et al., 2016), the central role of the ACC in affective and attentional pain processing (Garcia-Larrea et al., 2003) is certainly related to its multi-facet cognitive functionality and participation in domain-general control (Atlas, 2023; Wager et al., 2016).

The observation that the connectivity of the FM theta-alpha network was overall suppressed during mediation implies that meditative states inhibit the maintenance of pain representations and their communication across cortical regions, thus minimizing the role of pain information in the complex coordination of behaviour (Duprez et al., 2020). Notably, STM manifested a suppression of theta-alpha FM synchronization only during LKM pointing to (1) pronounced influence of positive emotional shifts and (2) the presence of strong interactions between emotional and fronto-medial cognitive control network in novice meditators. It will be interesting to explore whether this effect is specific to pain processing or also applies to other, non-nociceptive stimuli. Given the similar effects on P3b during LKM in STM, generalized effects of emotional activation on different cognitive systems can be suggested for novices. In contrast, in LTM, theta-alpha FM synchronization was reduced during FAM and OMM. Hence, the enhanced attention regulation imposed by these meditation forms was efficient to block the distribution and integration of nociceptive signals pointing to the presence of dominant interactions between attentional and medio-frontal integrative cognitive systems in LTM.

At the same time, the synchronization of FM beta networks was enhanced during FAM and OMM in LTM, indicating that fast frequency networks are specifically entrained during enhanced attention regulation in experienced meditators. Previously, beta synchronization has been associated with a top-down amplification of attended information (Buschman & Miller, 2007), or with mediating the link between top-down attentional selection and awareness (Driver & Mattingley, 1998; Fries, 2015; Gaillard et al., 2009; Gross et al., 2004; Yordanova et al., 2017). Also, medial frontal beta activity has been associated with monitoring of conflicts and subsequent behavioural adjustments and adaptation (Zavala et al., 2018). Thus, although the suppressed connectedness of the theta-alpha FM hub reveals a restricted integration of nociceptive signals during meditation in LTM, the co-existing increase of MF beta connectivity in FAM and OMM implies a simultaneous intensification of pain event monitoring possibly related to raised awareness. This observation is of special interest as it points to a capacity of expert meditators to employ attention mechanisms in order to segregate precisely information processing streams: While they could minimize the integration of negative pain representations as reflected by suppressed FM theta-alpha synchronization, high-frequency networks were simultaneously recruited to support advanced awareness of sensory pain signals.

### 4.9. Attentional fronto-parietal networks

Analyses of FP PLV during pain processing in different meditation states revealed that effects of meditation forms on theta-alpha FP synchronization were only present in experienced meditators. Specifically, a lateralized asymmetric pattern appeared for dorsal attention networks, as reflected by state-specific modulations of P3-F3 and P4-F4 electrode pairs (Fig. 3F). They were characterized by a left-hemisphere increase/right-hemisphere decrease during FAM, and an opposite right-hemisphere increase/left-hemisphere decrease during OMM. This hemispheric asymmetry may result from the different roles that the right and left hemispheres play for the association between executive attention and awareness (Posner & Rothbart, 1998). Specifically, it has been suggested that attention networks in the right hemisphere are engaged in top-down controlled sensory awareness, while attention networks in the left hemisphere are responsible for a voluntary motor control (Rounis et al., 2007; Rushworth et al., 1997, 2001, 2003). This account is consistent with the key processes required by OMM including open monitoring and enhanced awareness of ongoing events (Malinowski, 2013), which might have evoked a greater activation of right-hemisphere attentional networks. An additional explanation for the FP hemispheric asymmetry observed here may be linked with a right-lateralized superimposed influence from the ventral attention network controlling attention re-orienting to distracting, novel or salient events is right-lateralized (Chica et al., 2011, 2013; Corbetta & Shulman, 2002; Vossel et al., 2014). It can be suggested that in OMM, in contrast to FAM, the need to monitor consciously all external and internal events by avoiding at the same time any distraction to a specific signal, produced a greater activation of the right-hemisphere ventral network. In contrast, since the ventral network has been found to be suppressed during guided focused attention to restrict distraction and protect goal-driven targets (Shulman et al., 2003, 2007; Todd et al., 2005), which was the intended state in FAM, a strong suppression in the right hemisphere may be responsible for the left-hemisphere dominance during FAM. It is to be emphasized that meditation states did not produce significant changes in the synchronization of FP networks in novice meditators suggesting that these networks had limited role for distinctive pain processing during mediation in this group.

## 5. Overview and conclusions

Considering the findings of the present study as a whole shows that the maintenance of different meditation states does not affect bottom-up sensory processing of pain information, but affects significantly top-down neural mechanisms in state- and trait-dependent ways.

(1) In novice meditators, top-down influences are essentially guided by intended shifts to positive emotional reflection, with sustained attention having little effects. The meditation-raised emotional state mainly limits the distribution and cognitive/affective appraisal of nociceptive signals, which reduces the subjective experience of pain (intensity, aversion, and identification).
(2) In contrast, the effects of meditation on neural pain processing in experts are critically guided by advanced control of internal attention. They include (a) a pro-active decrease in the excitability of somatosensory areas, (b) a highly specialized reactivity of the fronto-medial cognitive network suppressing the communication and integration of nociceptive signals while simultaneously supporting raised awareness, (c) a highly specialized lateralization of attention networks reinforcing different types of attentional focus. Although these neural modulations are not accompanied by changes in subjective pain sensation, they reveal a fine-grained tuning and functional segregation of cognitive control and attention networks during meditation. These mechanisms in long-term mediators are possibly associated with a neuroplastic reorganization of large scale cognitive and pain processing networks that is a result of long-term engagement with meditation and has not yet developed in less experienced meditators.

The present results may have specific therapeutic value as they point to a role contemplative states that induce positive emotions and attitudes may play: Generating positive emotional states may be a useful approach for subjective pain relief in general population. Only experienced meditators can additionally alter specific pain processes during meditation by engaging advanced and highly-segregated functionality of cognitive control and attention networks. Thus, long-term attention training using specific meditation practices may provide a relevant additional approach in practical pain management.

Regarding the scientific understanding of pain, the combined consideration of meditation state and trait effects may have relevant implications for existing models of pain processes in meditation. In light of our previous results that meditation expertise essentially alters bottom-up neural mechanisms of pain (Yordanova et al., 2025), the model according to which meditation is associated with lower signalling in somatosensory areas and diminished processing of the sensory pain component due to a downregulation of ascending nociceptive signals (Jinich-Diamant et al., 2020; Zeidan & Vago, 2016; Zeidan et al., 2011) appears to refer to long-term trait effects of meditation. In view of the current results, the model proposing that meditation alters pain sensation by separating the sensory experience of pain from the corresponding evaluation of unpleasantness/affect (Gard et al., 2012; Grant et al., 2011; Perlman et al., 2010; Zeidan & Vago, 2016; Zorn et al., 2020, 2021), appears to apply mostly to phasic changes of brain states associated with different meditation forms.

## Acknowledgments

We would like to express our gratitude and appreciation to the monks, nuns and novices of Amaravati and Santacittarama Buddhist Monasteries for their outstanding dedication and participation in our study. This work has been supported by a grant from BIAL Foundation (Portugal) for the project “Advancements on the aware mind-brain: New insights about the neural correlates of meditation states and traits”, and by the National Research Fund by the Ministry of Education and Science, Sofia, Bulgaria (Project KP-06-N33/11/2019).

## Data availability

The datasets used and analysed during the current study are available from the corresponding author on reasonable request.

## References

[1] Apkarian, A. V., Bushnell, M. C., Treede, R. D., & Zubieta, J. K. (2005). Human brain mechanisms of pain perception and regulation in health and disease. Eur. J. Pain, 9, 463–484. 10.1016/j.ejpain.2004.11.001

[2] Atlas, L. Y. (2023). How instructions, learning, and expectations shape pain and neurobiological responses. Annu. Rev. Neurosci., 46, 167–189. 10.1146/annurev-neuro-101822-122427

[3] Babiloni, C., Babiloni, F., Carducci, F., Cincotti, F., Rosciarelli, F., Arendt-Nielsen, L., Chen, A. C., & Rossini, P.M. (2002). Human brain oscillatory activity phase-locked to painful electrical stimulations: a multi-channel EEG study. Hum. Brain Mapp., 15, 112–123. 10.1002/hbm.10013

[4] Babiloni, C., Brancucci, A., Babiloni, F., Capotosto, P., Carducci, F., Cincotti, F., Arendt-Nielsen, L., Chen, A. C., & Rossini, P. M. (2003). Anticipatory cortical responses during the expectancy of a predictable painful stimulation. A high-resolution electroencephalography study. Eur. J. Neurosci., 18, 1692–1700. 10.1046/j.1460-9568.2003.02851.x

[5] Babiloni, C., Brancucci, A., Arendt-Nielsen, L., Babiloni, F., Capotosto, P., Carducci, F., Cincotti, F., Del Percio, C., Petrini, L., Rossini, P. M., & Chen, A. C. (2004). Attentional processes and cognitive performance during expectancy of painful galvanic stimulations: a high-resolution EEG study. Behav. Brain Res., 152, 137–147. 10.1016/j.bbr.2003.10.004

[6] Babiloni, C., Brancucci, A., Del Percio, C., Capotosto, P., Arendt-Nielsen, L., Chen, A. C., & Rossini, P. M. (2006). Anticipatory electroencephalography alpha rhythm predicts subjective perception of pain intensity. J. Pain, 7, 709–717. 10.1016/j.jpain.2006.03.005

[7] Bakhshani, N. M., Amirani, A., Amirifard, H., & Shahrakipoor, M. (2016). The effectiveness of mindfulness-based stress reduction on perceived pain intensity and quality of life in patients with chronic headache. Glob. J. Health Sci., 8, 47326. 10.5539/gjhs.v8n4p142

[8] Benjamini, Y., & Hochberg, Y. (1995). Controlling the false discovery rate: a practical and powerful approach to multiple testing. R. Stat. Soc. B., 57, 289–300.

[9] Bingel, U., Lorenz, J., Glauche, V., Knab, R., Gläscher, J., Weiller, C., & Büchel, C. (2004). Somatotopic organization of human somatosensory cortices for pain: a single trial fMRI study. NeuroImage, 23, 224–232. 10.1016/j.neuroimage.2004.05.021

[10] Bott, F. S., Nickel, M. M., Hohn, V. D., May, E. S., Gil Ávila, C., Tiemann, L., Gross, J., & Ploner, M. (2023). Local brain oscillations and interregional connectivity differentially serve sensory and expectation effects on pain. Sci. Adv., 9, eadd7572. 10.1126/sciadv.add7572

[11] Brefczynski-Lewis, J. A., Lutz, A., Schaefer, H. S., Levinson, D. B., & Davidson, R. J. (2007). Neural correlates of attentional expertise in long-term meditation practitioners. Proc. Natl. Acad. Sci. USA, 104, 11483–11488. 10.1073/pnas.0606552104

[12] Bressler, S. L., & Menon, V. (2010). Large-scale brain networks in cognition: emerging methods and principles. Trends Cogn. Sci., 14, 277–290. 10.1016/j.tics.2010.04.004

[13] Brown, C. A., & Jones, A. K. (2010). Meditation experience predicts less negative appraisal of pain: Electrophysiological evidence for the involvement of anticipatory neural responses. Pain, 150, 428–438. 10.1016/j.pain.2010.04.017

[14] Buschman, T. J., & Miller, E. K. (2007). Top-down versus bottom-up control of attention in the prefrontal and posterior parietal cortices. Science, 315(5820), 1860–1864. 10.1126/science.1138071

[15] Cahn, B. R., & Polich, J. (2006). Meditation states and traits: EEG, ERP, and neuroimaging studies. Psychol. Bull., 132, 180–211. 10.1037/0033-2909.132.2.180

[16] Cavanagh, J. F., & Frank, M. J. (2014). Frontal theta as a mechanism for cognitive control. Trends Cogn. Sci., 18, 414–421. 10.1016/j.tics.2014.04.012

[17] Cavanagh, J. F., Cohen, M. X., & Allen, J. J. (2009). Prelude to and resolution of an error: EEG phase synchrony reveals cognitive control dynamics during action monitoring. J. Neurosci., 29, 98–105. 10.1523/JNEUROSCI.4137-08.2009

[18] Chica, A. B., Bartolomeo, P., & Valero-Cabré, A. (2011). Dorsal and ventral parietal contributions to spatial orienting in the human brain. J. Neurosci., 31, 8143–8149. 10.1523/jneurosci.5463-10.2010

[19] Chica, A. B., Paz-Alonso, P. M., Valero-Cabré, A., & Bartolomeo, P. (2013). Neural bases of the interactions between spatial attention and conscious perception. Cereb. Cortex, 23, 1269–1279. 10.1093/cercor/bhs087

[20] Chiesa, A., & Malinowski, P. (2011). Mindfulness-based approaches: are they all the same? J. Clin. Psychol., 67, 404–424. 10.1002/jclp.20776

[21] Cohen, M. X., & Donner, T. H. (2013). Midfrontal conflict-related theta-band power reflects neural oscillations that predict behavior. J. Neurophysiol., 110, 2752–2763. 10.1152/jn.00479.2013

[22] Cohen, M. X., & van Gaal, S. (2013). Dynamic interactions between large-scale brain networks predict behavioral adaptation after perceptual errors. Cereb. Cortex, 23, 1061–1072. 10.1093/cercor/bhs069

[23] Cohen, M. X. (2011). Error-related medial frontal theta activity predicts cingulate-related structural connectivity. NeuroImage, 55, 1373–1383. 10.1016/j.neuroimage.2010.12.072

[24] Cohen, M. X. (2014).A neural microcircuit for cognitive conflict detection and signaling. Trends Neurosci., 37, 480–490. 10.1016/j.tins.2014.06.004

[25] Corbetta, M., & Shulman, G. L. (2002). Control of goal-directed and stimulus-driven attention in the brain. Nat. Rev. Neurosci., 3, 201–215. 10.1038/nrn755

[26] Corbetta, M., & Shulman, G. L. (2011). Spatial neglect and attention networks. Ann. Rev. Neurosci., 34, 569–599. 10.1146/annurev-neuro-061010-113731

[27] Dahl, C. J., Lutz, A., & Davidson, R. J. (2015). Reconstructing and deconstructing the self in three families of meditation. Trends Cogn. Sci., 19, 1–9. 10.1016/j.tics.2015.07.001

[28] Daitch, A. L., Sharma, M., Roland, J. L., Astafiev, S. V., Bundy, D. T., Gaona, C. M., Snyder, A. Z., Shulman, G. L., Leuthardt, E. C., & Corbetta, M. (2013). Frequency-specific mechanism links human brain networks for spatial attention. Proc. Natl. Acad. Sci. USA, 110, 19585–19590. 10.1073/pnas.1307947110

[29] Davidson, R. J., & Dahl, C. J. (2017). Varieties of contemplative practice. JAMA Psychiatry, 74, 121–123. 10.1001/jamapsychiatry.2016.3469

[30] Dehaene, S., & Changeux, J. P. (2011). Experimental and theoretical approaches to conscious processing. Neuron, 70, 200–227. 10.1016/j.neuron.2011.03.018

[31] Dehaene, S., Kerszberg, M., & Changeux, J. P. (1998). A neuronal model of a global workspace in effortful cognitive tasks. Proc. Natl. Acad. Sci. USA, 95, 14529–14534. 10.1073/pnas.95.24.14529

[32] Della Penna, S., Torquati, K., Pizzella, V., Babiloni, C., Franciotti, R., Rossini, P. M., & Romani, G. L. (2004). Temporal dynamics of alpha and beta rhythms in human SI and SII after galvanic median nerve stimulation. A MEG study. NeuroImage, 22, 1438–1446. 10.1016/j.neuroimage.2004.03.045

[33] Dockstader, C., Cheyne, D., & Tannock, R. (2010). Cortical dynamics of selective attention to somatosensory events. NeuroImage, 49, 1777–1785. 10.1016/j.neuroimage.2009.09.035

[34] Driver, J., & Mattingley, J. B. (1998). Parietal neglect and visual awareness. Nat. Neurosci., 1, 17–22. 10.1038/217

[35] Duprez, J., Gulbinaite, R., & Cohen, M. X. (2020). Midfrontal theta phase coordinates behaviorally relevant brain computations during cognitive control. NeuroImage, 207, 116340. 10.1016/j.neuroimage.2019.116340

[36] Egner, T., Monti, J. M., Trittschuh, E. H., Wieneke, C. A., Hirsch, J., & Mesulam, M. M. (2008). Neural integration of top-down spatial and feature-based information in visual search. J. Neurosci., 28, 6141–6151. 10.1523/JNEUROSCI.1262-08.2008

[37] Fell, J., Axmacher, N., & Haupt, S. (2010). From alpha to gamma: electrophysiological correlates of meditation-related states of consciousness. Med. Hypotheses, 75, 218–224. https://doi.og/10.1016/j.mehy.2010.02.025

[38] Fox, K. C. R., Nijeboer, S., Dixon, M. L., Floman, J. L., Ellamil, M., Rumak, S. P., Sedlmeier, P., & Christoff, K. (2014). Is meditation associated with altered brain structure? A systematic review and meta-analysis of morphometric neuroimaging in meditation practitioners. Neurosci. Biobehav. Rev., 43, 48–73. 10.1016/j.neubiorev.2014.03.016

[39] Fries, P. (2015). Rhythms for cognition: Communication through coherence. Neuron, 88, 220–235. 10.1016/j.neuron.2015.09.034

[40] Fusco, G., Scandola, M., Feurra, M., Pavone, E. F., Rossi, S., & Aglioti, S. M. (2018). Midfrontal theta transcranial alternating current stimulation modulates behavioural adjustment after error execution. Eur. J. Neurosci., 48, 3159–3170. 10.1111/ejn.14174

[41] Fusco, G., Fusaro, M., & Aglioti, S. M. (2022). Midfrontal-occipital θ-tACS modulates cognitive conflicts related to bodily stimuli. Soc. Cogn. Affect. Neurosci., 17, 91–100. 10.1093/scan/nsaa125

[42] Gaillard, R., Dehaene, S., Adam, C., Clémenceau, S., Hasboun, D., Baulac, M., Cohen, L., & Naccache, L. (2009). Converging intracranial markers of conscious access. PLoS Biol., 7, e61. 10.1371/journal.pbio.1000061

[43] Garcia-Larrea, L., & Peyron, R. (2013). Pain matrices and neuropathic pain matrices: a review. Pain, 154, Suppl. 1, S29–S43. 10.1016/j.pain.2013.09.001

[44] Garcia-Larrea, L., Frot, M., & Valeriani, M. (2003). Brain generators of laser-evoked potentials: from dipoles to functional significance. Neurophysiol. Clin., 33, 279–292. 10.1016/j.neucli.2003.10.008

[45] Gard, T., Hölzel, B. K., Sack, A. T., Hempel, H., Lazar, S. W., Vaitl, D., & Ott, U. (2012). Pain attenuation through mindfulness is associated with decreased cognitive control and increased sensory processing in the brain. Cereb. Cortex, 22, 2692–2702. 10.1093/cercor/bhr352

[46] Grant, J. A., & Rainville, P. (2009). Pain sensitivity and analgesic effects of mindful states in Zen meditators: a cross-sectional study. Psychosom. Med., 71, 106–114. 10.1097/psy.0b013e31818f52ee

[47] Grant, J. A., Courtemanche, J., & Rainville, P. (2011). A non-elaborative mental stance and decoupling of executive and pain-related cortices predicts low pain sensitivity in Zen meditators. Pain, 152, 150–156. 10.1016/j.pain.2010.10.006

[48] Gross, J., Schmitz, F., Schnitzler, I., Kessler, K., Shapiro, K., Hommel, B., & Schnitzler A. (2004). Modulation of long-range neural synchrony reflects temporal limitations of visual attention in humans. Proc. Natl. Acad. Sci. USA, 101, 13050–13055. 10.1073/pnas.0404944101

[49] Grossman, P., Tiefenthaler-Gilmer, U., Raysz, A., & Kesper, U. (2007). Mindfulness training as an intervention for fibromyalgia: evidence of postintervention and 3-year follow-up benefits in well-being. Psychother. Psychosom., 76, 226. 10.1159/000101501

[50] Guidotti, R., D’Andrea, A., Basti, A., Raffone, A., Pizzella, V., & Marzetti, L. (2023). Long-term and meditation-specific modulations of brain connectivity revealed through multivariate pattern analysis. Brain Topogr., 36, 409–418. 10.1007/s10548-023-00950-3

[51] Hagiwara, K., Okamoto, T., Shigeto, H., Ogata, K., Somehara, Y., Matsushita, T., Kira, J., & Tobimatsu, S. (2010). Oscillatory gamma synchronization binds the primary and secondary somatosensory areas in humans. NeuroImage, 51, 412–420. 10.1016/j.neuroimage.2010.02.001

[52] Hasenkamp, W., Wilson-Mendenhall, C. D., Duncan, E., & Barsalou, L. W. (2012). Mind wandering and attention during focused meditation: A fine-grained temporal analysis of fluctuating cognitive states. NeuroImage, 59, 750–760. 10.1016/j.neuroimage.2011.07.008

[53] Hauck, M., Lorenz, J., & Engel, A. K. (2007). Attention to painful stimulation enhances gamma-band activity and synchronization in human sensorimotor cortex. J. Neurosci., 27, 9270–9277. 10.1523/jneurosci.2283-07.2007

[54] Hauck, M., Domnick, C., Lorenz, J., Gerloff, C., & Engel, A. K. (2015). Top-down and bottom-up modulation of pain induced oscillations. Front. Hum. Neurosci., 9, 375. 10.3389/fnhum.2015.00375

[55] Huneke, N. T., Brown, C. A., Burford, E., Watson, A., Trujillo-Barreto, N. J., El-Deredy, W., & Jones, A. K. (2013). Experimental placebo analgesia changes resting-state alpha oscillations. PLoS One, 8, e78278. 10.1371/journal.pone.0078278

[56] Isbel, B., & Summers, M. J. (2017). Distinguishing the cognitive processes of mindfulness: Developing a standardised mindfulness technique for use in longitudinal randomised control trials. Consc. Cogn., 52, 75–92. 10.1016/j.concog.2017.04.019

[57] Jinich-Diamant, A., Garland, E., Baumgartner, J., Gonzalez, N., Riegner, G., Birenbaum, J., Case, L., & Zeidan, F. (2020). Neurophysiological mechanisms supporting mindfulness meditation-based pain relief: an updated review. Curr. Pain Headache Rep., 24, 56. 10.1007/s11916-020-00890-8

[58] Jo, H. G., Malinowski, P., & Schmidt, S. (2017). Frontal theta dynamics during response conflict in long-term mindfulness meditators. Front. Hum. Neurosci., 11, 299. 10.3389/fnhum.2017.00299

[59] Kasai, Y., Sakakibara, T., Kyaw, T. A., Soe, Z. W., Han, Z. M., & Htwe, M. M. (2017). Psychological effects of meditation at a Buddhist monastery in Myanmar. J. Ment. Health, 26, 4–7. 10.3109/09638237.2015.1124405

[60] Klimesch, W., Sauseng, P., & Hanslmayr, S. (2007). EEG alpha oscillations: the inhibition-timing hypothesis. Brain Res. Rev., 53, 63–88. 10.1016/j.brainresrev.2006.06.003

[61] Koessler, L., Benhadid, A., Maillard, L., Vignal, J. P., Felblinger, J., Vespignani, H., & Braun, M. (2008). Automatic localization and labeling of EEG sensors (ALLES) in MRI volume. NeuroImage, 41, 914–923. 10.1016/j.neuroimage.2008.02.039

[62] Kolev, V., Beste, C., Falkenstein, M., & Yordanova, J. (2009). Error-related oscillations: Effects of aging on neural systems for behavioral monitoring. J. Psychophysiol., 23, 216–223. 10.1027/0269-8803.23.4.216

[63] Kolev, V., Falkenstein, M., & Yordanova, J. (2024). A distributed theta network of error generation and processing in aging. Cogn. Neurodyn., 18, 447–459. 10.1007/s11571-023-10018-4

[64] Kucyi, A., & Davis, K. D. (2015). The dynamic pain connectome. Trends Neurosci., 38, 86–95. 10.1016/j.tins.2014.11.006

[65] Lachaux, J. P., Rodriguez, E., Martinerie, J., & Varela, F. J. (1999). Measuring phase synchrony in brain signals. Hum. Brain Mapp., 8, 194–208. 10.1002/(SICI)1097-0193(1999)8:4%3C194::AID-HBM4%3E3.0.CO;2-C

[66] Legrain, V., Guérit, J. M., Bruyer, R., & Plaghki, L. (2002). Attentional modulation of the nociceptive processing into the human brain: selective spatial attention, probability of stimulus occurrence, and target detection effects on laser evoked potentials. Pain, 99, 21–39. 10.1016/s0304-3959(02)00051-9

[67] Legrain, V., Bruyer, R., Guérit, J. M., & Plaghki, L. (2005). Involuntary orientation of attention to unattended deviant nociceptive stimuli is modulated by concomitant visual task difficulty. Evidence from laser evoked potentials. Clin. Neurophysiol., 116, 2165–2174. 10.1016/j.clinph.2005.05.019

[68] Legrain, V., Damme, S. V., Eccleston, C., Davis, K. D., Seminowicz, D. A., & Crombez, G. (2009). A neurocognitive model of attention to pain: behavioral and neuroimaging evidence. Pain, 144, 230–232. 10.1016/j.pain.2009.03.020

[69] Legrain, V., Crombez, G., & Mouraux, A. (2011). Controlling attention to nociceptive stimuli with working memory. PLoS One, 6, e20926. 10.1371/journal.pone.0020926

[70] Legrain, V., Mancini, F., Sambo, C. F., Torta, D. M., Ronga, I., & Valentini, E. (2012). Cognitive aspects of nociception and pain: bridging neurophysiology with cognitive psychology. Neurophysiol. Clin., 42, 325–336. 10.1016/j.neucli.2012.06.003

[71] Legrain, V., Crombez, G., Plaghki, L., & Mouraux, A. (2013). Shielding cognition from nociception with working memory. Cortex, 49, 1922–1934. 10.1016/j.cortex.2012.08.014

[72] Lu, C., Moliadze, V., & Nees, F. (2023). Dynamic processes of mindfulness-based alterations in pain perception. Front. Neurosci., 17, 1253559. 10.3389/fnins.2023.1253559

[73] Lutz, A., Slagter, H. A., Dunne, J. D., & Davidson, R. J. (2008). Attention regulation and monitoring in meditation. Trends Cogn. Sci., 12, 163–169. 10.1016/j.tics.2008.01.005

[74] Lutz, A., Jha, A. P., Dunne, J. D., & Saron, C. D. (2015). Investigating the phenomenological matrix of mindfulness-related practices from a neurocognitive perspective. Am. Psychol., 70, 632–658. 10.1037/a0039585

[75] Makeig, S., Bell, A. J., Jung, T. -P., Ghahremani, D., & Sejnowski, T. J. (1997). Blind separation of auditory event-related brain responses into independent components. Proc. Natl Acad. Sci. USA, 94, 10979–10984. 10.1073/pnas.94.20.10979

[76] Malinowski, P. (2013). Neural mechanisms of attentional control in mindfulness meditation. Front. Neurosci., 7, 8. 10.3389/fnins.2013.00008

[77] Mallat, S. (1999). A Wavelet Tour of Signal Processing, 2nd ed. (Academic Press, San Diego, CA).

[78] Menon, V. (2013). Developmental pathways to functional brain networks: emerging principles. Trends Cogn. Sci., 17, 627–640. 10.1016/j.tics.2013.09.015

[79] Morone, N. E., Greco, C. M., & Weiner, D. K. (2008). Mindfulness meditation for the treatment of chronic low back pain in older adults: a randomized controlled pilot study. Pain, 134, 310–319. 10.1016/j.pain.2007.04.038

[80] Nickel, M. M., Tiemann, L., Hohn, V. D., May, E. S., Gil Ávila, C., Eippert, F., & Ploner, M. (2022). Temporal-spectral signaling of sensory information and expectations in the cerebral processing of pain. Proc. Natl. Acad. Sci. USA, 119, e2116616119. 10.1073/pnas.2116616119

[81] Nicolardi, V., Simione, L., Scaringi, D., Malinowski, P., Yordanova, J., Kolev, V., Mauro, F., Giommi, F., Barendregt, H. P., Aglioti, S. M., & Raffone, A. (2022). The two arrows of pain: mechanisms of pain related to meditation and mental states of aversion and identification. Mindfulness, 15, 753–774. 10.1007/s12671-021-01797-0

[82] Nielsen, L., & Kaszniak, A. W. (2006). Awareness of subtle emotional feelings: a comparison of long-term meditators and nonmeditators. Emotion, 6, 392–405. 10.1037/1528-3542.6.3.392

[83] Nigbur, R., Cohen, M. X., Ridderinkhof, K. R., & Stürmer, B. (2012). Theta dynamics reveal domain-specific control over stimulus and response conflict. J. Cogn. Neurosci., 24, 1264–1274. 10.1162/jocn_a_00128

[84] Nunez, P. L., Srinivasan, R., Westdorp, A. F., Wijesinghe, R. S., Tucker, D. M., Silberstein, R. B., & Cadusch, P. J. (1997). EEG coherency. I: Statistics, reference electrode, volume conduction, Laplacians, cortical imaging, and interpretation at multiple scales. Electroencephalogr. Clin. Neurophysiol., 103, 499–515. 10.1016/s0013-4694(97)00066-7

[85] Peerdeman, K. J., van Laarhoven, A. I. M., Keij, S. M., Vase, L., Rovers, M. M., Peters, M. L., & Evers, A. W. M. (2016). Relieving patients’ pain with expectation interventions: a meta-analysis. Pain, 157, 1179–1191. 10.1097/j.pain.0000000000000540

[86] Peng, W., Babiloni, C., Mao, Y., & Hu, Y. (2015). Subjective pain perception mediated by alpha rhythms. Biol. Psychol., 109, 141–50. 10.1016/j.biopsycho.2015.05.004

[87] Perlman, D. M., Salomons, T. V., Davidson, R. J., & Lutz, A. (2010). Differential effects on pain intensity and unpleasantness of two meditation practices. Emotion, 10, 65. 10.1037/a0018440

[88] Perrin, F., Pernier, J., Bertrand, O., & Echallier, J. F. (1989). Spherical splines for scalp potential and current density mapping. Electroencephalogr. Clin. Neurophysiol., 72, 184–187. 10.1016/0013-4694(89)90180-6

[89] Peyron, R., Garcia-Larrea, L., Gregoire, M. C., Costes, N., Convers, P., Lavenne, F., Mauguiere, F., Michel, D., & Laurent, B. (1999). Haemodynamic brain responses to acute pain in humans: sensory and attentional networks. Brain, 122, 1765–1779. 10.1093/brain/122.9.1765

[90] Peyron, R., Laurent, B., & Garcia-Larrea, L. (2000). Functional imaging of brain responses to pain. A review and meta-analysis. Neurophysiol. Clin., 30, 263–288. 10.1016/s0987-7053(00)00227-6

[91] Pfurtscheller, G., & Lopes da Silva, F. H. (1999). Event-related EEG/MEG synchronization and desynchronization: basic principles. Clin. Neurophysiol., 110, 1842–1857. 10.1016/s1388-2457(99)00141-8

[92] Ploner, M., Gross, J., Timmermann, L., Pollok, B., & Schnitzler, A. (2006a). Oscillatory activity reflects the excitability of the human somatosensory system. NeuroImage, 32, 1231–1236. 10.1016/j.neuroimage.2006.06.004

[93] Ploner, M., Gross, J., Timmermann, L., Pollok, B., & Schnitzler, A. (2006b). Pain suppresses spontaneous brain rhythms. Cereb. Cortex, 16, 537–540. 10.1093/cercor/bhj001

[94] Ploner, M., Sorg, C., & Gross, J. (2017). Brain rhythms of pain. Trends Cogn. Sci., 21, 100–110. 10.1016/j.tics.2016.12.001

[95] Polich, J. (2007). Updating P300: an integrative theory of P3a and P3b. Clin. Neurophysiol., 118, 2128–2148. 10.1016/j.clinph.2007.04.019

[96] Posner, M., & Rothbart, M. K. (1998). Attention, self-regulation and consciousness. Phil. Trans. R. Soc. Lond. B, 353, 1915–1927. 10.1098/rstb.1998.0344

[97] Raffone, A., & Srinivasan, N. (2009). An adaptive workspace hypothesis about the neural correlates of consciousness: insights from neuroscience and meditation studies. Progr. Brain Res., 176, 161–180. 10.1016/S0079-6123(09)17620-3

[98] Raffone, A., Srinivasan, N., & Barendregt, H. P. (2014). Attention, consciousness and mindfulness in meditation. Psychology of Meditation, pp. 147–166 (Nova Science Publishers).

[99] Raffone, A., Marzetti, L., Del Gratta, C., Perrucci, M. G., Romani, G. L., & Pizzella, V. (2019). Toward a brain theory of meditation. Progr. Brain Res., 244, 207–232. 10.1016/bs.pbr.2018.10.028

[100] Rees, G. (2013). Neural correlates of consciousness. Ann. NY Acad. Sci., 1296, 4–10. 10.1111/nyas.12257

[101] Rottschy, C., Langner, R., Dogan, I., Reetz, K., Laird, A. R., Schulz, J. B., Fox, P. T., & Eickhoff, S. B. (2012). Modelling neural correlates of working memory: a coordinate-based meta-analysis. NeuroImage, 60, 830–846. 10.1016/j.neuroimage.2011.11.050

[102] Rottschy, C., Caspers, S., Roski, C., Reetz, K., Dogan, I., Schulz, J. B., Zilles, K., Laird, A. R., Fox, P. T., & Eickhoff, S. B. (2013). Differentiated parietal connectivity of frontal regions for "what" and "where" memory. Brain Struct. Funct., 218, 1551–1567. 10.1007/s00429-012-0476-4

[103] Rounis, E., Yarrow, K., & Rothwell, J. C. (2007). Effects of rTMS conditioning over the fronto-parietal network on motor versus visual attention. J. Cogn. Neurosci., 19, 513–524. 10.1162/jocn.2007.19.3.513

[104] Rushworth, M. F., Nixon, P. D., Renowden, S., Wade, D. T., & Passingham, R. E. (1997). The left parietal cortex and motor attention. Neuropsychologia, 35, 1261–1273. 10.1016/s0028-3932(97)00050-x

[105] Rushworth, M. F., Krams, M., & Passingham, R. E. (2001). The attentional role of the left parietal cortex: the distinct lateralization and localization of motor attention in the human brain. J. Cogn. Neurosci., 13, 698–710. 10.1162/089892901750363244

[106] Rushworth, M. F., Johansen-Berg, H., Göbel, S. M., & Devlin, J. T. (2003). The left parietal and premotor cortices: motor attention and selection. NeuroImage, 20 (Suppl. 1), S89–100. 10.1016/j.neuroimage.2003.09.011

[107] Sadaghiani, S., & Kleinschmidt, A. (2016). Brain networks and α-oscillations: Structural and functional foundations of cognitive control. Trends Cogn. Sci., 20, 805–817. 10.1016/j.tics.2016.09.004

[108] Säterö, P., Klingenstierna, U., Karlsson, T., & Olausson, B. (2000). Pain threshold measurements with cutaneous argon laser, comparing a forced choice and a method of limits. Prog. Neuropsychopharmacol. Biol. Psychiatry, 24, 397–407. 10.1016/s0278-5846(99)00107-4

[109] Schmidt, S., Grossman, P., Schwarzer, B., Jena, S., Naumann, J., & Walach, H. (2011). Treating fibromyalgia with mindfulness-based stress reduction: results from a 3-armed randomized controlled trial. Pain, 152, 361–369. 10.1016/j.pain.2010.10.043

[110] Scrivener, C. L., & Reader, A. T. (2022). Variability of EEG electrode positions and their underlying brain regions: visualizing gel artifacts from a simultaneous EEG-fMRI dataset. Brain Behav., 12, e2476. 10.1002/brb3.2476

[111] Shulman, G. L., McAvoy, M. P., Cowan, M. C., Astafiev, S. V., Tansy, A. P., d’Avossa, G., & Corbetta, M. (2003). Quantitative analysis of attention and detection signals during visual search. J. Neurophysiol., 90, 3384–3397. 10.1152/jn.00343.2003

[112] Shulman, G. L., Astafiev, S. V., McAvoy, M. P., d’Avossa, G., & Corbetta, M. (2007). Right TPJ deactivation during visual search: functional significance and support for a filter hypothesis. Cereb. Cortex, 17, 2625–2633. 10.1093/cercor/bhl170

[113] Singh, A., Patel, D., Li, A., Hu, L., Zhang, Q., Liu, Y., Guo, X., Robinson, E., Martinez, E., Doan, L., Rudy, B., Chen, Z. S., & Wang, J. (2020). Mapping cortical integration of sensory and affective pain pathways. Curr. Biol., 30, 1703–1715.e5. 10.1016/j.cub.2020.02.091

[114] Strube, A., Rose, M., Fazeli, S., & Buchel, C. (2021). Spatial and spectral characteristics of expectations and prediction errors in pain and thermoception. eLife, 10, e62809. 10.7554/eLife.62809

[115] Su, I. W., Wu, F. W., Liang, K. C., Cheng, K. Y., Hsieh, S. T., Sun, W. Z., & Chou, T. L. (2016). Pain perception can be modulated by mindfulness training: a resting-state fMRI study. Front. Hum. Neurosci., 10, 570. 10.3389/fnhum.2016.00570

[116] Tallon-Baudry, C., Bertrand, O., Delpuech, C., & Permier, J. (1997). Oscillatory gamma-band (30-70 Hz) activity induced by a visual search task in humans. J. Neurosci., 17, 722–734. 10.1523/JNEUROSCI.17-02-00722.1997

[117] Tang, Y. Y., Hölzel, B. K., & Posner, M. I. (2015). The neuroscience of mindfulness meditation. Nat. Rev. Neurosci., 16, 213–225. 10.1038/nrn3916

[118] Tiemann, L., May, E. S., Postorino, M., Schulz, E., Nickel, M. M., Bingel, U., & Ploner, M. (2015). Differential neurophysiological correlates of bottom-up and top-down modulations of pain. Pain, 156, 289–296. 10.1097/01.j.pain.0000460309.94442.44

[119] Todd, J. J., Fougnie, D., & Marois, R. (2005). Visual short-term memory load suppresses temporo-parietal junction activity and induces inattentional blindness. Psychol. Sci., 16, 965–972. 10.1111/j.1467-9280.2005.01645.x

[120] Vago, D. R., & Silbersweig, D. A. (2012). Self-awareness, self-regulation, and self-transcendence (S-ART): a framework for understanding the neurobiological mechanisms of mindfulness. Front. Hum. Neurosci., 6, 296. 10.3389/fnhum.2012.00296

[121] Valentini, E., Nicolardi, V., & Aglioti, S. M. (2017). Painful engrams: Oscillatory correlates of working memory for phasic nociceptive laser stimuli. Brain Cogn., 115, 21–32. 10.1016/j.bandc.2017.03.009

[122] van Ede, F., Jensen, O., & Maris, E. (2010). Tactile expectation modulates pre-stimulus beta-band oscillations in human sensorimotor cortex. NeuroImage, 51, 867–876. 10.1016/j.neuroimage.2010.02.053

[123] van Ede, F., de Lange, F., Jensen, O., & Maris, E. (2011). Orienting attention to an upcoming tactile event involves a spatially and temporally specific modulation of sensorimotor alpha- and beta-band oscillations. J. Neurosci., 31, 2016–2024. 10.1523/JNEUROSCI.5630-10.2011

[124] Vossel, S., Geng, J. J., & Fink, G. R. (2014). Dorsal and ventral attention systems: Distinct neural circuits but collaborative roles. Neuroscientist, 20, 150–159. 10.1177/1073858413494269

[125] Wager, T. D., Atlas, L. Y., Botvinick, M. M., Chang, L. J., Coghill, R. C., Davis, K. D., Iannetti, G. D., Poldrack, R. A., Shackman, A. J., & Yarkoni, T. (2016). Pain in the ACC? Proc. Natl. Acad. Sci. USA, 113, E2474–2475. 10.1073/pnas.1600282113

[126] Wiech, K. (2016). Deconstructing the sensation of pain: The influence of cognitive processes on pain perception. Science, 354, 584–587. 10.1126/science.aaf8934

[127] Yordanova, J., Falkenstein, M., Hohnsbein, J., & Kolev, V. (2004). Parallel systems of error processing in the brain. NeuroImage, 22, 590–602. 10.1016/j.neuroimage.2004.01.040

[128] Yordanova, J., Kolev, V., Verleger, R., Heide, W., Grumbt, M., & Schürmann, M. (2017). Synchronization of fronto-parietal beta and theta networks as a signature of visual awareness in neglect. NeuroImage, 146, 341–354. 10.1016/j.neuroimage.2016.11.013

[129] Yordanova, J., Falkenstein, M., & Kolev, V. (2020). Aging-related changes in motor response-related theta activity. Int. J. Psychophysiol., 153, 95–106. 10.1016/j.ijpsycho.2020.03.005

[130] Yordanova, J., Kolev, V., Mauro, F., Nicolardi, V., Simione, L., Calabrese, L., Malinowski, P., & Raffone, A. (2020). Common and distinct lateralised patterns of neural coupling during focused attention, open monitoring and loving kindness meditation. Sci. Rep., 10, 7430. 10.1038/s41598-020-64324-6

[131] Yordanova, J., Kolev, V., Nicolardi, V., Simione, L., Mauro, F., Garberi, P., Raffone, A., & Malinowski, P. (2021). Attentional and cognitive monitoring brain networks in long-term meditators depend on meditation states and expertise. Sci. Rep., 11, 4909. 10.1038/s41598-021-84325-3

[132] Yordanova, J., Falkenstein, M., & Kolev, V. (2024). Aging alters functional connectivity of motor theta networks during sensorimotor reactions. Clin. Neurophysiol., 158, 137–148. 10.1016/j.clinph.2023.12.132

[133] Yordanova, J., Nicolardi, V., Malinowski, P., Simione, L., Aglioti, S. M., Raffone, A., & Kolev, V. (2025). EEG oscillations reveal neuroplastic changes in pain processing associated with long-term meditation. Sci. Rep., 15, 10604. 10.1038/s41598-025-94223-7

[134] Zavala, B., Jang, A., Trotta, M., Lungu, C. I., Brown, P., & Zaghloul, K. A. (2018). Cognitive control involves theta power within trials and beta power across trials in the prefrontal-subthalamic network. Brain, 141, 3361–3376. 10.1093/brain/awy266

[135] Zeidan, F., & Vago, D. R. (2016). Mindfulness meditation–based pain relief: a mechanistic account. Ann. NY Acad. Sci., 1373, 114–127. 10.1111/NYAS.13153

[136] Zeidan, F., Martucci, K. T., Kraft, R. A., Gordon, N. S., McHaffie, J. G., & Coghill, R. C. (2011). Brain mechanisms supporting the modulation of pain by mindfulness meditation. J. Neurosci., 31, 5540–5548. 10.1523/JNEUROSCI.5791-10.2011

[137] Zeidan, F., Emerson, N. M., Farris, S. R., Ray, J. N., Jung, Y., McHaffie, J. G., & Coghill, R. C. (2015). Mindfulness meditation-based pain relief employs different neural mechanisms than placebo and sham mindfulness meditation-induced analgesia. J. Neurosci., 35, 15307–15325. 10.1523/jneurosci.2542-15.2015

[138] Zorn, J., Abdoun, O., Bouet, R., & Lutz, A. (2020). Mindfulness meditation is related to sensory-affective uncoupling of pain in trained novice and expert practitioners. Eur. J. Pain, 24, 1301–1313. 10.1002/ejp.1576

[139] Zorn, J., Abdoun, O., Sonié, S., & Lutz, A. (2021). Cognitive defusion is a core cognitive mechanism for the sensory-affective uncoupling of pain during mindfulness meditation. Psychosom. Med., 83, 566–578. 10.1097/PSY.0000000000000938

